# Speciation of the *Mycobacterium tuberculosis* complex using mass spectrometry: a proof-of-concept

**DOI:** 10.1101/2021.02.04.429756

**Authors:** Simon Robinne, Jamal Saad, Madjid Morsli, Hamidou Zelika Harouna, Fatah Tazerart, Michel Drancourt, Sophie Alexandra Baron

## Abstract

Mycobacteria that form the *Mycobacterium tuberculosis* complex are responsible for deadly tuberculosis in animals and patients. Identification of these pathogens at the species level is of primary importance for treatment and source tracing, and currently relies on DNA analysis, including whole genome sequencing (WGS), which takes a whole day. In this study, we report on the unprecedented identification of the *M. tuberculosis* complex species using matrix-assisted laser desorption ionization-time of flight mass spectrometry (MALDI-TOF-MS), with WGS as the comparative gold standard. In a first step, an optimised peptide extraction applied to 24 isolates otherwise identified in three of the 11 *M. tuberculosis* complex species by WGS, yielded 94 MALDI-TOF spectra which clustered according to WGS identification. In a second step, 70/74 (95%) other isolates were correctly identified at the species level by this clustering method. This study is the first to report a MALDI-TOF-MS method of identification of *M. tuberculosis* complex mycobacteria at the species level and is easily implantable in clinical microbiology laboratories.

## INTRODUCTION

Mycobacteria that form the *Mycobacterium tuberculosis* complex are responsible for tuberculosis, a deadly infection that is a major public health concern around the world (1). The *M. tuberculosis* complex is composed of 11 different species: *Mycobacterium tuberculosis, Mycobacterium africanum, Mycobacterium orygis, Mycobacterium bovis, Mycobacterium microti, Mycobacterium canettii, Mycobacterium caprae, Mycobacterium pinnipedii, Mycobacterium suricattae, Mycobacterium dassie*, and *Mycobacterium mungi* (also related as ecotypes) (2). In our clinical microbiology practice however, *M. tuberculosis stricto sensu*, along with *M. bovis* and its derivative, Bacillus Calmette-Guérin (BCG), and *M. africanum*, are the sole *M. tuberculosis* complex species that we have encountered in 10 years.

The identification of *M. tuberculosis* isolates at the species level is a concern for the immediate medical management of patients as, for example, *M. bovis* and BCG are two species acknowledged to be pyrazinamide-resistant (3). It is also of concern in terms of tracing the source in order to prevent additional cases, in the case of zoonotic *M. bovis* tuberculosis for example (4).

While real-time PCR detection of the *M. tuberculosis* complex directly in clinical samples is now widely used for the routine screening of tuberculosis, this technique does not allow for a specific identification within the complex (5). In routine clinical microbiology, the identification of the *M. tuberculosis* complex at the species level is achieved by DNA sequence investigations and, at best, by whole genome sequencing (WGS) of the colony (5). These performing techniques, however, require specific equipment that may not be available at the point-of-care, with results taking a few days to become available (Saad J., Baron S., Amrane S., Gallou J., Brieu N. and Drancourt M., submitted for publication).

Matrix-assisted laser desorption ionization-time of flight mass spectrometry (MALDI-TOF-MS), a widely implanted technique for the routine identification of bacteria in clinical microbiology laboratories as a rapid and lower cost method for identifying bacterial species (6, 7), is also routinely used for the identification of mycobacteria (8, 9). However, MALDI-TOF-MS suffers from the same limitation as real-time PCR, i.e., it is impossible to differentiate *M. tuberculosis* complex species based on similar peptide profiles measurable with currently available MALDI-TOF-MS protocols. The results is that it is currently impossible to specify tuberculosis species based on this method (10).

In this study, we investigated whether clustering MALDI-TOF-MS spectra obtained from *M. tuberculosis* complex colonies using principal component analysis (PCA) could achieve species identification, by cross-referencing the data retrieved from genome sequencing. PCA clustering has already proven to be a mathematical technique which can be widely applied to MALDI-TOF-MS datasets, and is able to discriminate between groups of isolates based on their peptide profile (11).

## MATERIALS AND METHODS

### Mycobacterial isolates

A total of 105 clinical isolates cultured in the BSL3 laboratory of the IHU Méditerranée Infection, Marseille, France (N=85) and the Laboratoire National de Référence des IST/VIH et de la Tuberculose, Niamey, Niger (N=20) were included in this study. This collection of isolates has previously been investigated, in part for accurate identification by WGS, here used as the gold standard (Saad J., Baron S., Amrane S., Gallou J., Brieu N. and Drancourt M, submitted for publication).

In a first phase, 24 isolates from the IHU Méditerranée Infection, identified at the species level by WGS in the *M. tuberculosis* complex (*M. tuberculosis, M. africanum, M. bovis*), were defined as reference strains for the construction of a standard PCA cluster of MALDI-TOF-MS spectra. These isolates were selected to be representative of the diversity of species and genetic lineages encountered at the IHU Méditerannée Infection laboratory.

In a second phase, 74 other isolates from the IHU Méditerranée Infection (Marseille, France) and the Laboratoire National de Référence des IST/VIH et de la Tuberculose (Niamey, Niger), identified at the species level by WGS in the *M. tuberculosis* complex (*M. tuberculosis, M. africanum, M. bovis*), were defined as our test group, in order to evaluate the reliability of the identification of the standard cluster previously generated. Finally, seven isolates from the IHU Méditerranée Infection, identified at the species level by WGS outside the *M. tuberculosis* complex (two isolates of *Mycobacterium abscessus*, two *Mycobacterium avium, Mycobacterium chelonae, Mycobacterium gilvum, Mycobacterium persicum*) were used as negative controls of our PCA clustering method. All isolates were cultured on TransBK m4 agar medium (Culture-Top, Les Ulys, France), a culture medium derived from Middlebrook 7H10 agar medium at 37°C under a 5% CO2 atmosphere for a minimum of seven days. Total DNA extraction and WGS were performed as previously described (Saad J., Baron S., Amrane S., Gallou J., Brieu N. and Drancourt M, submitted for publication).

### MALDI-TOF-MS analysis

Between one and three 1-μL plastic loops harvested from each isolate were mixed with 300 μL of high purity liquid chromatography (HPLC)-grade water into a 1.5-mL tube (Sarstedt, Nümbrecht, Germany). Samples were then heated to 100°C for 30 minutes in order to inactivate mycobacteria (12). After cooling, 900 μL of ethanol was added, vortexed for one minute and centrifuged for two minutes at 16,060×g, and the supernatant was eliminated. The centrifugation step was repeated twice, and the pellet was dried at room temperature for several minutes. A spatula-full of glass beads for cell disruption (diameter: 0.5mm; Scientific Industries Inc., Bohemia, USA) and 20 μL of acetonitrile were added, then samples were stirred using a vortex device for one minute. 20 μL of a solution of 70% formic acid was then added and agitated for five seconds. Samples were then agitated using a FastPrep-24^™^ device (MP Biomedicals, #6004500) for five cycles of 20 seconds, with a resting time of five seconds between each cycle, then centrifuged for two minutes at 16,060xg. Four replicates of 1.5μL supernatant were deposited on a MALDI target plate (MSP 96 target polished steel, Bruker Daltonics, Bremen, Germany) and left to dry at room temperature for one minute. Matrix solution consisting of 1.5μL of a saturated □-cyano 4-hydroxycinnamic acid (CHCA) (Sigma-Aldrich, Taufkirchen, Germany) in TFA 5% acetonitrile (1:1) was deposited on each spot and left to dry for five minutes at room temperature. Two spots consisting of 1.5-μL CHCA matrix only were incorporated as negative controls and two spots consisting of 1.5-μL 1/10 diluted solution of Bruker Bacterial Standard Test, an *Escherichia coli* extract spiked with two high molecular weight proteins (Bruker Daltonics), mixed with 1.5-μL of CHCA matrix were deposited on every plate, as positive controls. All the solvents used were MS grade. Mass spectra were obtained on a MicroFlex^™^ (Bruker Daltonics) MALDI-TOF MS. Spectra were recorded in the mass range from 2,000 to 20,000 Da.

### Spectra processing

Spectra were processed using the Biotyper Compass Explorer v4.1 (Bruker Daltonics). Each spectrum was compared with the Mycobacteria bead method Library v.3.0 (Bruker Daltonics). PCA spectra dendrograms were performed using the default parameter set (distance value measured by correlation model with average linkage algorithm, number of clusters maximum defined at four). Dendrogram distances were built by the software on an arbitrary unit based on dissimilarities between spectra, with a range from 0 to 1.2.

After the definition of the list of spectra used for the building of the standard cluster, each quadrupled of spectra obtained on the same isolate from the test group were added to the standard list in order to perform a PCA clustering. 81 dendrograms were thus generated, one for each isolate of the negative and test group. On each dendrogram, the distance separating MALDI-TOF-MS spectra from the tested isolate by spectra from the closest isolate in the reference group was measured. Species from the closest isolate in the reference group were also noted.

## RESULTS

### Speciation of mycobacteria based on database match

MALDI-TOF-MS spectra presenting a low intensity (under 2,000 a.u.) were removed from the study, providing a total number of 411 MALDI-TOF-MS spectra of *Mycobacterium* species investigated (Tables 1–2), with a mean of 3.91 spectra per isolate. Identification of spectra by comparison with the MALDI Biotyper Compass IVD Library v9.0 and the Mycobacteria bead method Library v.3.0 (Bruker Daltonics) yielded 69% correct identifications of *Mycobacterium* species with a comparison score above 2.00; and this value rose to 93% using a comparison score above 1.80.

**Table 1.** Mycobacterial isolates and associated spectra used for building standard PCA clusters

| Species |  | Number of isolates | Number of spectra |
| --- | --- | --- | --- |
| *Mycobacterium africanum* |  | 2 | 8 |
| *Mycobacterium bovis* |  | 3 | 12 |
| *Mycobacterium bovis* (BCG) |  | 2 | 8 |
| *Mycobacterium tuberculosis* | Lineage 1 (Indo-oceanic) | 3* | 12* |
|  | Lineage 2 (East-Asia) | 3 | 12 |
|  | Lineage 3 (East-African-Indian) | 3 | 11 |
|  | Lineage 4 (Euro-American) | 8 | 31 |
| TOTAL |  | 24* | 94* |

\*: four spectra from one isolate of *M. tuberculosis* (lineage 1.1.1, Indo-Oceanic) were removed from the reference cluster dataset as it clusterised incorrectly.

**Table 2.** Observations on individual PCA dendrogram generated on spectra from mycobacteria isolates of the test group

| Isolates species | Number of isolates | Number of spectra | Mean distance of clusterisation | Standard deviation of distance of clusterisation | Consistency of clusterisation with closest species |
| --- | --- | --- | --- | --- | --- |
| **Negative control:** |  |  |  |  |  |
| *Mycobacterium abscessus* | 2 | 8 | 0.827 | NA | NA |
| *Mycobacterium avium* | 2 | 8 | 0.938 | NA | NA |
| *Mycobacterium chelonae* | 1 | 4 | 0.928 | NA | NA |
| *Mycobacterium gilvum* | 1 | 4 | 0.995 | NA | NA |
| *Mycobacterium persicum* | 1 | 4 | 0.795 | NA | NA |
| TOTAL Negative controls | 7 | 28 | 0.892 | 0.094 | NA |
| **Test group:** |  |  |  |  |  |
| *Mycobacterium africanum* | 1 | 4 | 0.333 | NA | 0% |
| *Mycobacterium bovis* | 3 | 12 | 0.458 | 0.067 | 66.7% |
| *Mycobacterium bovis (BCG)* | 1 | 4 | 0.750 | NA | 100% |
| *Mycobacterium tuberculosis* | 69 | 269 | 0.387 | 0.167 | 97.1% |
| TOTAL Test group | 74 | 289 | 0.398 | 0.167 | 94.6% |

### Reference cluster

Figure 1 shows the standard PCA dendrogram built with the 90 spectra (Table 1) obtained from *M. tuberculosis* complex isolates defined as reference groups. One *M. tuberculosis* isolate was erroneously included in group 3 (*M. bovis*/ *M. africanum*) and was thus excluded from the reference cluster. Interestingly, this isolate had been identified by WGS as being a *M. tuberculosis* lineage 1.1.1 (Indo-Oceanic), the lineage of *M. tuberculosis* most closely related to *M. africanum* and *M. bovis* (13). Accordingly, the reference cluster used for further identification of isolates consisted of 23 isolates. The maximum average distance among spectra obtained from replicates of the same bacterial isolate was calculated at 0.420 ± 0.202. As shown in Figure 1, the standard cluster presented three main groups here referred to as group 1, 2 and 3. Group 1 presents a top node of separation measured at 0.810, and a clusterisation of spectra coherent with species identification, being composed of spectra obtained on 11 different isolates of *M. tuberculosis*. Group 2 (top node of separation measured at 0.568) presents a coherent clustering with species identification of spectra from five different isolates of *M. tuberculosis*. Group 3 (top node of separation measured at 0.897) presents clustering of spectra obtained from *M. africanum* (2), *M. bovis* (1) and *M. bovis* BCG isolates (2). No consistent clustering according to *M. tuberculosis* lineages was observed in clusters 1 and 2 (Figure 2). Reference cluster spectra were deposited on the Méditerranée Infection website (https://www.mediterranee-infection.com/acces-ressources/base-de-donnees/urms-data-base/#link_acc-25-26-d) and are publicly accessible using doi 10.35081/mbbf-rm88 (https://urldefense.com/v3/__https://www.mediterranee-infection.com/wp-content/uploads/2021/01/Speciation-of-mycobacteria-by-MALDI-TOF-Reference-cluster-dataset.zip__;!!JQ5agg!I6GLl-a6ZbO9zQRBdgO1YspBNlOOliEkFyqzKjjwmo0TsExFDQu_FUN5rlC07DPa8jpx$).

**Figure 1:**
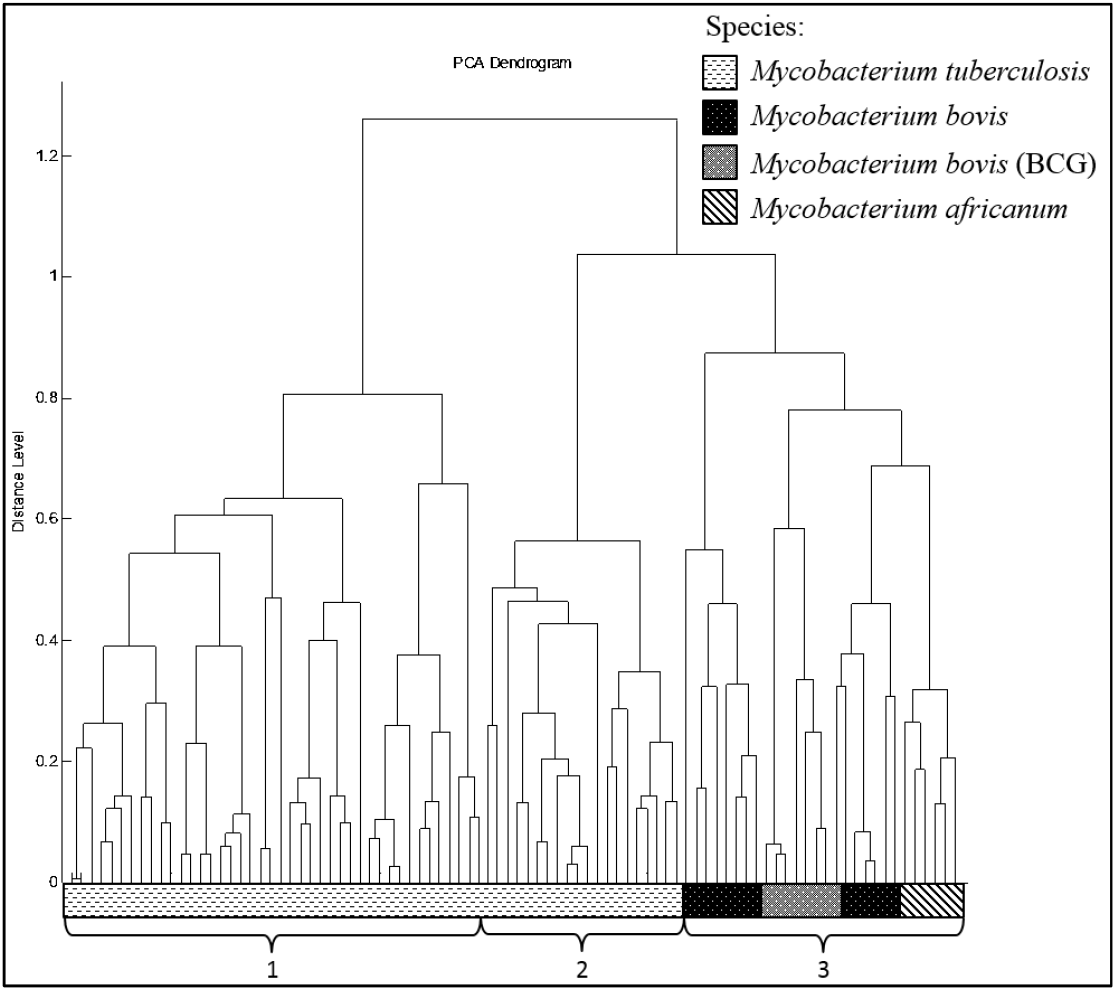
Standard PCA dendrogram generated with spectra from mycobacteria isolates of the standard group

**Figure 2:**
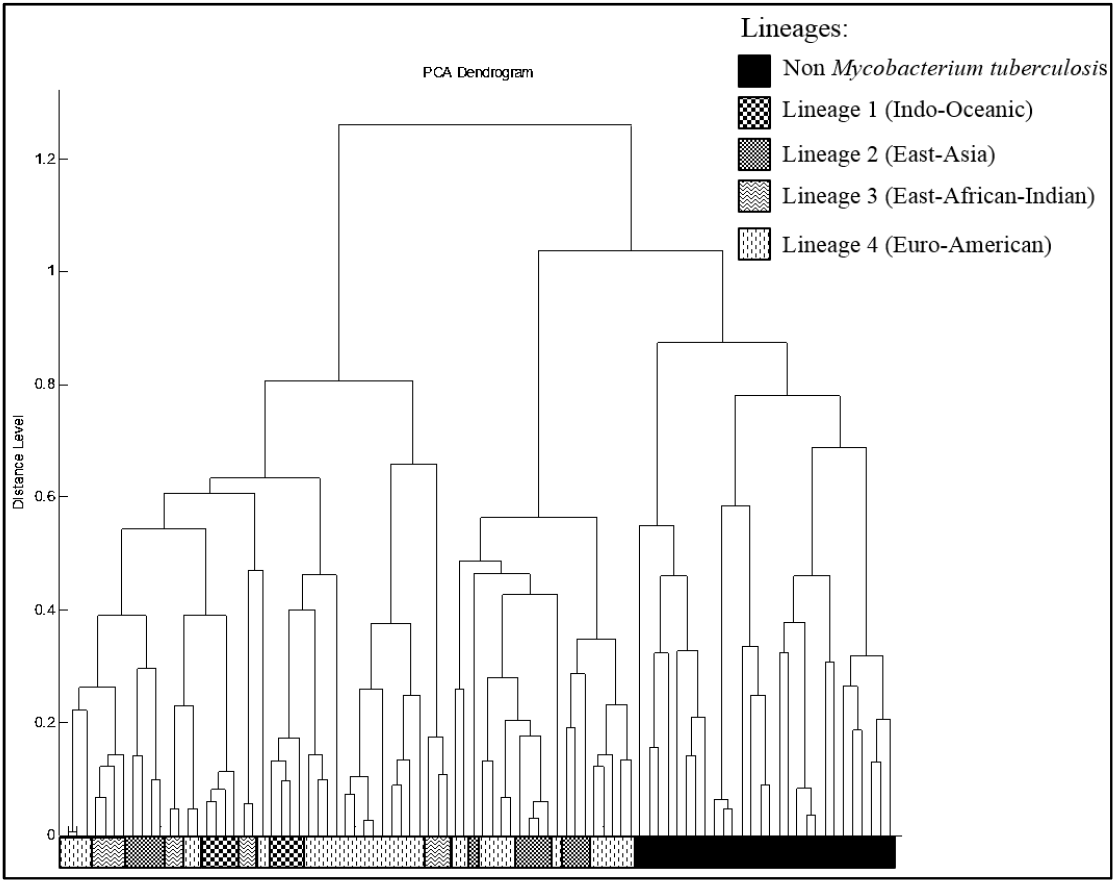
Observation of clusterisation of *Mycobacterium tuberculosis* lineages among standard PCA dendrograms

### Identification of mycobacteria using reference clustering

Observations of 81 dendrograms generated with spectra added to the spectra from isolates of the test group are summarised in Table 2. Overall, 77/81 (95.1%) isolates were correctly identified at the species level (using WGS as the gold standard for identification), including 70/74 (94.6%) isolates identified in the *M. tuberculosis* complex and 7/7 isolates identified outside the *M. tuberculosis* complex. In contrast, one *M. bovis* isolate was misidentified as *M. tuberculosis*,one *M. africanum* isolate as *M. bovis*, and two *M. tuberculosis* isolates (lineages 2.2.1 and 1.1.2) as *M. bovis*.

## DISCUSSION

In this study, we report the unprecedented identification of *M. tuberculosis* complex species using the MALDI-TOF-MS approach. Data here reported were validated by the fact that positive and negative controls deposited on each MALDI-TOF-MS target plate respectively correctly identified *E. coli* (score above 2.00) and no identifications (score under 1.60) against selected database; MALDI-TOF-MS spectra were repeated over at least four repeats; negative controls of the PCA clustering method consisting of 7 isolates of mycobacteria species not included in the *M. tuberculosis* complex (*M. abscessus*, *M. avium*, *M. chelonae*, *M. gilvum, M. persicum*) presented spectra with clusterisation distance significantly higher than spectra from isolates of the test groups in the *M. tuberculosis* complex (0.892±0.094 versus 0.398±0.167; p value =2.39E-7; Student’s t-test); all identifications were validated using WGS as the gold standard and spectra from the reference group were deposited in a publicly available database.

Here, spectra analysis not only consisted in matching with a database, as reported for most previous studies dealing with MALDI-TOF-MS identification of mycobacteria as a whole and *M. tuberculosis* complex in particular (8, 9), but rather a clustering approach consisting of a PCA against a standard dataset of spectra; similar to the approach previously reported by O’Connor *et al*. on combined summary spectra (10). Consequently, we are reporting for the first time to the best of our knowledge, the identification of *M. tuberculosis* complex isolates at the species level, starting from colonies. More precisely, this involves the PCA dendrogram carried out on the 90 spectra of the standard group discriminated *M. tuberculosis* isolates (groups 1 and 2) and *non-M. tuberculosis* isolates (group 3). Coherent clustering of spectra was observed for 70 isolates among 74 constituting test group (94.6%), involving a reliable accuracy of clusterisation with the standard cluster among spectra of the same mycobacteria species inside the *M. tuberculosis* complex. However, of the 74 isolates in the test group, only five clinical isolates were non-tuberculosis mycobacteria, and accuracy of clustering among these five isolates was only 60%. Two inconsistent clusterings consisted in one case of one *M. africanum* isolate that clustered with spectra from *M. bovis* in group 3. As it was observed in group 3 of the standard cluster that spectra from *M. africanum* and *M. bovis* presented close clustering, this could indicate that the PCA clustering method is not suitable for distinguishing these two species, but does make it possible to distinguish between them and *M. tuberculosis*. Indeed, O’Connor *et al*. have already observed that clusterisation of the MALDI-TOF dataset acquired from isolates of the *M. tuberculosis* complex presented distinct grouping of two on three *M. bovis* (BCG) isolates compared to other *M. tuberculosis* isolates, which is coherent with what was observed on the dataset used here (10).

Indeed, rapid identification of *M. tuberculosis* complex isolates at the species level is of medical interest, giving a preliminary information regarding the potential antibiotic susceptibility pattern, as *M. bovis* and *M. caprae* are intrinsically resistant to pyrazinamide (14) and potential sources of contamination such as *M. bovis, M. caprae* and *M. pinnipedii* are three species acknowledged as being responsible for zoonotic tuberculosis (4). In this study however, the lack of consistency of clustering according to *M. tuberculosis* lineage among groups 1 and 2 indicated that the PCA clustering method reported here did not discriminate the *M. tuberculosis* lineage.

In conclusion, we report on the proof-of-concept that PCA clustering of MALDI-TOF-MS spectra of mycobacteria can be used as a first line method of identification of colonies in the *M. tuberculosis* complex; presenting a reasonable positive predictive value of identification of 94.6%. The method here reported may be refined by increasing the database and may be extended to mycobacteria grown in broth, as all data here presented were acquired from colonies grown on a solid culture medium. The PCA clustering reported here is a more accurate tool for identifying dissimilarities inside groups of closely related spectra than the current identification by comparison with a database. As such, it could be used to investigate ways of discriminating isolates within other bacteria complexes that are still unsolved by MALDI-TOF MS identification.

## ACKNOWLEDGEMENTS

The authors are very grateful to Gilles Liotaud, Simon Pinchemel and Amaël Fadlane (Assistance Publique-Hôpitaux de Marseille, BSL3 laboratory and Strain Collection of the Rickettsia Unit) for their technical work on mycobacteria culture. MM would like to thank Ghiles Grine for the organisation of genome sequencing. Finally, the authors would like to acknowledge the Head of the Laboratoire National de Référence des IST/VIH et de la Tuberculose, in Niamey, Niger.

## SUPPLEMENTARY MATERIAL

The data presented in this study are openly available on the Méditerranée Infection website (https://www.mediterranee-infection.com/acces-ressources/base-de-donnees/urms-data-base/#link_acc-25-26-d) at [doi: 10.35081/mbbf-rm88].

